# A VEGF reaction-diffusion mechanism that selects variable densities of endothelial tip cells

**DOI:** 10.1101/624999

**Authors:** W. Bedell, A. D. Stroock

## Abstract

The patterned differentiation of endothelial cells into tip and stalk cells represents an important step in the process of angiogenic sprouting. Vascular biologists hypothesize that changes in the density and overall structure of the vasculature can be traced in part to changes in the number of tip cells selected in the endothelium prior to sprout formation. However, the dominant hypotheses for tip cell selection invoke lateral inhibition via Notch; this juxtacrine mechanism predicts that a fixed fraction of endothelial cells become tip cells through a pattern-forming instability. Here, we present and analyze a hypothetical mechanism for tip cell selection that is based on endothelial competition for diffusible vascular endothelial growth factor (VEGF); this mechanism predicts that variable densities of tip cells emerge depending on the local (paracrine) production rate of VEGF. First, we hypothesize a network of VEGF signaling and trafficking based on previous experimental findings that could allow internalization of VEGF to occur with positive feedback. We formalize the hypothesis into a set of nonlinear ordinary differential equations and perform linear stability analysis to elucidate a general criterion for tip cell pattern formation under the mechanism. We use numerical integration to explore the nonlinear dynamics and final steady-states of tip cell patterns under this mechanism; the observed density of tip cells can be tuned from 10% to 84%. We conclude with proposals of future experiments and computational studies to explore how competitive consumption of diffusible VEGF may play a role in determining vascular structure.

**Statement of Significance:** The patterned differentiation of endothelial cells into tip and stalk cells represents an important step in the process of blood vessel growth. Vascular biologists hypothesize that changes in the density and overall structure of the vasculature can be traced in part to changes in the number of tip cells selected during angiogenesis. However, the dominant hypotheses for tip cell selection predict that a locally fixed fraction of endothelial cells become tip cells following stimulation by vascular endothelial growth factor (VEGF). Here, we present and analyze a hypothetical mechanism for tip cell selection based on endothelial competition for diffusible VEGF; this mechanism predicts that variable densities of tip cells emerge depending on the local production rate of VEGF.

## Introduction

Sprouting angiogenesis builds new blood vessels in early development, wound healing, and tumor growth (1). The process requires a population of endothelial cells to perform repeated cycles of collective differentiation and sprouting (Fig. 1) (2). Continued advances of angiogenesis-related therapies (3), methods for fabricating artificial tissues (4–6), and our fundamental understanding of vascular development (2) depend on the ability to explain and predict the multicellular processes governing angiogenesis.

Many early attempts to control blood vessel growth looked for an on-off switch – molecular or biophysical stimuli that could halt or induce the formation of angiogenic sprouts (7). However, commonly used angiogenesis inhibitors, such as the antibodies targeting vascular endothelial growth factor (VEGF), have been most successful at reducing the density of vascular sprouting, rather than eliminating it altogether (8–11). To explain how the density of sprouts is governed *in vivo*, many researchers have pointed to changes in the density of tip cell selection prior to sprouting (Fig. 1) (12, 13).

Upon stimulation by VEGF from nearby tissues, quiescent endothelial cells (ECs) spontaneously differentiate into a pattern of cells with the competency to be become either migratory tip cells or proliferative stalk cells (12). Given additional time, these tip- or stalk-competent cells undergo morphological change to become the tips or stalks of new angiogenic sprouts (14). A number of investigators have suggested that the balance of tip cell selection – the number of tip cells selected and their levels of activity – impacts the structure of the vasculature produced by angiogenesis, in particular, whether the vascular network is pathologically dense or sparse (13–16).

A dominant hypothesis in this field contends that tip cells are selected via a mechanism of lateral inhibition: nascent tip cells express high levels of the Delta ligand, which activates Notch in adjacent ECs to repress Delta expression and tip cell characteristics (12, 14, 17). Previous computational studies have shown that lateral inhibition, while robust in its ability to enable tip cell differentiation, is limited to forming fixed patterns of alternating tip and stalk cells when simulated in regular grids; this pattern represents a “checkerboard” of 50% tip cells in square grids (18, 19) or a pattern that can flip between either 33% or 66% tip cells in hexagonal lattices (20). This juxtacrine mechanism that only selects a fixed fraction (or flips between two fixed fractions) of tip cells is not compatible with observations of variable sprouting density (10, 16, 21).

**Figure 1:**
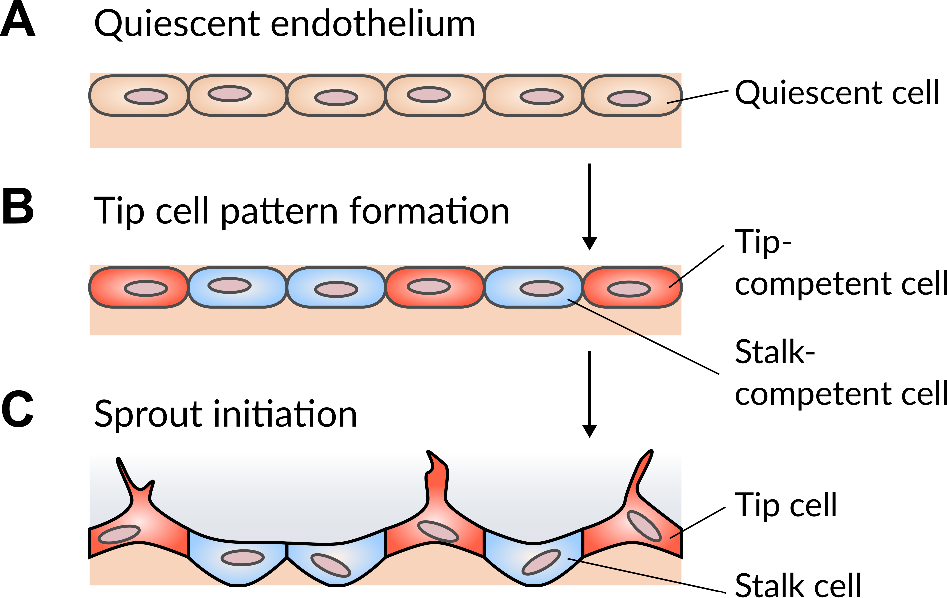
Overview of tip cell pattern formation. (A-B) Quiescent endothelial cells (A) are stimulated by angiogenic factors to differentiate into tip- (red) and stalk-competent (blue) cells (B). (C) Endothelial cells that become tip-competent during pattern formation become the tips of angiogenic sprouts during sprout initiation.

Mechanisms that predict variable tip cell density while remaining dependent on lateral inhibition via Notch have been proposed, but they face some challenges. A previous study showed that by considering filopodia that may carry Delta ligand to distant neighbors, lateral inhibition could create multicellular patterns less dense than the previously-mentioned “checkerboards” (22); while endothelial tip cells do express filopodia, it remains to be shown that these filopodia are essential for tip cell selection (23).

Another route to predicting variable numbers of tip cells selected via lateral inhibition during angiogenesis is to spatially restrict tip cell pattern formation to “patches” within the endothelium. In one previous example, Boareto et al. (24) proposed and simulated a mechanism that included both lateral inhibition and lateral induction (the convergence of neighboring cell phenotypes) via Notch. The final steady states in the model by Boareto et al. were composed of patches of approximately 50% tips cells surrounded by uniformly differentiated non-tip cell endothelium – the global numbers of tip cells varied according to the total area of tip cell-containing patches relative to non-patterned endothelium (24). In a previous computational study, Bentley et al. (18) proposed pathological angiogenesis as a result of “patchy” tip cell formation in a manner that also relied on Notch-based lateral inhibition. Recent experiments (19) have lent some support to aspects of this hypothesis by showing that chronically high VEGF can result in patches of high Delta expression, but experimental support for the existence of “checkerboard” patterns that locally select ∼50% of endothelial cells in non-pathological angiogenesis has yet to emerge.

Alternative mechanisms of vascular patterning without juxtacrine signaling have been explored. For example, early computational studies of angiogenesis, which did not invoke tip cell selection, had no role for juxtacrine Notch signaling (25, 26). Rather, VEGF would stimulate endothelial cells to divide and migrate towards its source: sprouts would initiate at whichever endothelial cell was exposed to the highest concentration of VEGF. As the endothelial cells migrated and consumed the VEGF, branching vascular structures could emerge. In contrast, most computational studies explicitly incorporating tip cell selection have assumed that the concentration of extracellular VEGF is constant across the endothelium or is a fixed function of position (18, 24, 27). In this study, we explore a hybrid of these two hypotheses: we attribute the sites of sprout initiation to patterning of tip and stalk phenotypes, with this patterning being regulated by the reaction-diffusion of VEGF. Relative to existing approaches, we invoke neither cell migration nor juxtacrine signaling.

Fig. 2A depicts a hypothetical mechanism of tip cell selection based on competition for available VEGF in the local extracellular space. Briefly, an endothelial cell must internalize VEGF at a high rate to become a tip cell, depleting the local concentration of VEGF and thus preventing neighboring cells from become tip cells. The fraction of tip cells selected depends on the rate at which VEGF is introduced from adjacent tissues: a stronger flux of VEGF can sustain more tip cells in the final pattern. From a multicellular perspective, this hypothesis, that limitations of VEGF control tip cell density, is consistent with previous experiment observations showing that VEGF receptors (VEGFR2) displayed by adjacent neurons act as strong sinks for VEGF that limit local formation of endothelial tip cells (28, 29).

We hypothesize that the signaling network responsible for this reaction-diffusion patterning (Fig. 2B) begins with the binding of VEGF to VEGFR2 displayed by endothelial cells themselves. Next, VEGF-VEGFR2 complexes move into dynamin-dependent vesicles; previous experiments have shown that movement of VEGF-VEGFR2 into such vesicles is required for VEGF-stimulated activation of Akt (a.k.a. protein kinase B) (34, 37). Active Akt has been shown to rapidly increase the production of nitric oxide (NO) in endothelial cells by activating endothelial nitric oxide synthase (eNOS) (31). NO enhances the ability of dynamin to form vesicles (32) and is enriched in, and important for the formation of, endothelial tip cells (33). Together, these observations suggest that intracellular VEGF-VEGFR2 provides signals to increase the activity of dynamin through Akt and NO; successful tip cells form the most vesicles, have the strongest signaling from internalized VEGF-VEGFR2 complexes, and produce the most NO (31–33). Finally, the action of Akt drives commitment to the tip phenotype. Our hypothesis is consistent with experiments showing that endothelial internalization of VEGF is required for full VEGF-VEGFR2 activation and tip cell formation (34–36).

In this study, we translate the hypothesis of Fig. 2 into a system of nonlinear ordinary differential equations (ODEs). We track the diffusion, internalization, and signaling of VEGF to help understand whether the hypothesized mechanism can select variable densities of endothelial tip cells. From these equations, we extract an exact stability criterion for the onset of spontaneous differentiation and we numerically simulate the full dynamics of tip cell pattern formation. We show that the VEGF reaction-diffusion mechanism is capable of generating tip cell patterns with densities that vary with increasing or decreasing VEGF production rates, in a manner consistent with previous experimental observations that vascular structure is responsive to VEGF concentration – given that, at steady state, we expect local concentration to vary linearly with flux (8–11).

**Figure 2:**
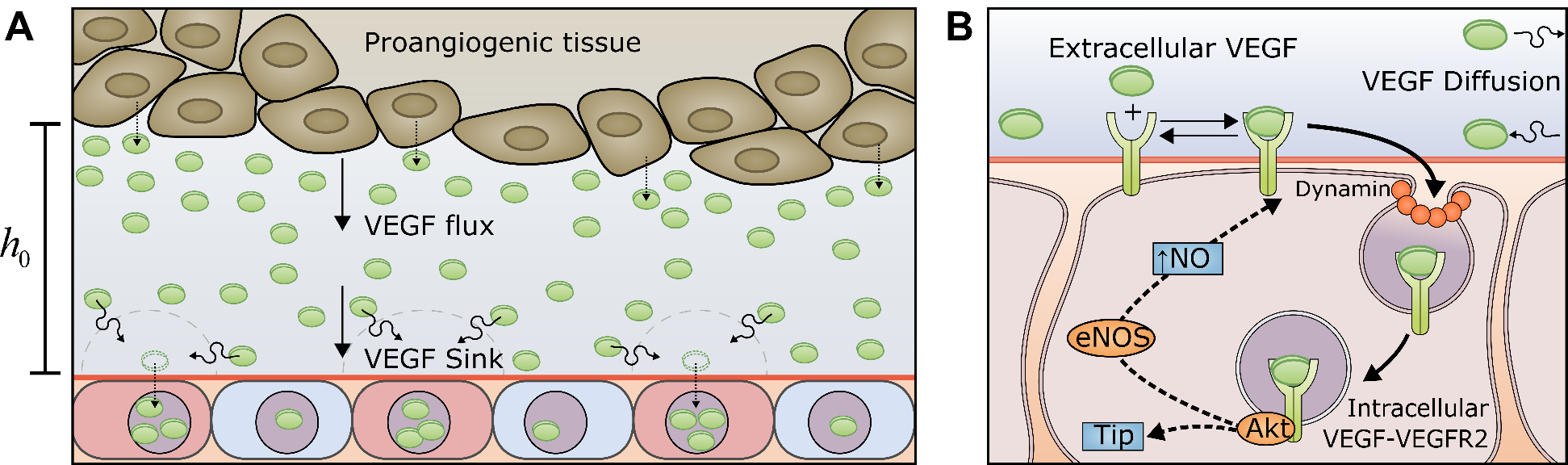
Hypothetical VEGF reaction-diffusion mechanism for tip cell selection. Proangiogenic tissues continuously produce a flux of VEGF across the extracellular space (with assumed height *h*_0_[μm], not shown to scale) towards endothelial cells (ECs). When exposed to high concentrations of VEGF, ECs spontaneously differentiate into tip- (red) and stalk- (blue) competent cells (30). We hypothesize that nascent tip cells have a high rate of VEGF consumption, creating a local sink in VEGF concentration that drives an increased flux in VEGF towards their cell surface. (B) We propose a network of VEGF trafficking and signaling that could enable the hypothesized reaction-diffusion mechanism: endosomal VEGF signaling may accelerate the rate of VEGF capture in dynamin-dependent vesicles (with Akt, eNOs, and NO performing intermediate signaling steps) causing cells with high levels of endosomal VEGF to consume VEGF more quickly (31–33) and adopt the tip phenotype (34–36).

## Model formulation

The primary goal of this study is to understand if the mechanism hypothesized in Fig. 2 exhibits an instability that leads to spontaneous tip cell pattern formation when simulated on a lattice of initially uniform endothelial cells. Tip cell selection can occur within a wall of a mature blood vessel (Fig. 3A, bottom left) or in a growing vascular plexus (Fig. 3A, top left); in both cases, the endothelium forms a 2D structure. We approximate the endothelium as a lattice of square-shaped compartments (Fig. 3A, right) with length and width *l* [cm], each representing an endothelial cell. Our approximation of the endothelium as a square lattice (see Supplemental S1) allows us to employ a connectivity matrix (****M****) to calculate the average concentration of extracellular VEGF in neighboring cells (i.e., 〈*V*〉 _*i*_ = *M*_*ij*_*V*_*i*_, where 〈*V*〉_*i*_ is the average concentration of extracellular VEGF in fluid volumes neighboring cell *i*); following Othmer & Scriven (38), this construction also allows us to perform a linear stability analysis to predict whether a pattern-forming instability can be created by the hypothesized mechanism for the coupled dynamics of signaling, mass transport, and gene expression.

Fig. 3B presents a cross-section of the local environment considered in the model: within the fluid volume associated with each cell *i* within the endothelium, extracellular VEGF (*V*_*i*_[mol/cm^3^]) is introduced as a constant flux 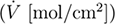 from nearby tissues. The height of the fluid volume (*h*_0_ [cm]) defines the local capacitance of the system and thus determines how sensitive the concentration of extracellular VEGF is to changes in the rate of VEGF internalization. In this fluid volume, we approximate the concentration of VEGF (*V*_*i*_ [mol/cm^3^]) as uniform in the direction normal to the cell membrane (*z*), but that it diffuses along gradients in the lateral directions (*x*, *y*) with a diffusivity *D*_*V*_ [cm^2^/s]. This approximation is reasonable for the geometry considered in this study, with a lateral dimension of cells of 20 μm and a height of the adjacent fluid volume of *h*_0_ = 10 μm (see Supplemental S1). Extracellular VEGF may reversibly bind to VEGFR2 at the endothelial cell surface. We hypothesize that internalization of VEGFR2 occurs continuously via a dynamin-dependent process (34, 37); when the internalized VEGFR2 is bound to extracellular VEGF, extra-cellular VEGF is captured into intracellular VEGF-VEGFR2 complexes (*I*_*i*_ [mol/cm^2^]) in the approximately two-dimensional cell. We assume that the intracellular VEGF-VEGFR2 complexes degrade via a first-order process. We further hypothesize that signaling from intracellular VEGF-VEGFR2 complexes accelerates the rate of VEGFR2 internalization, and by extension, increases the maximum rate at which extracellular VEGF can be internalized (as discussed below).

**Figure 3:**
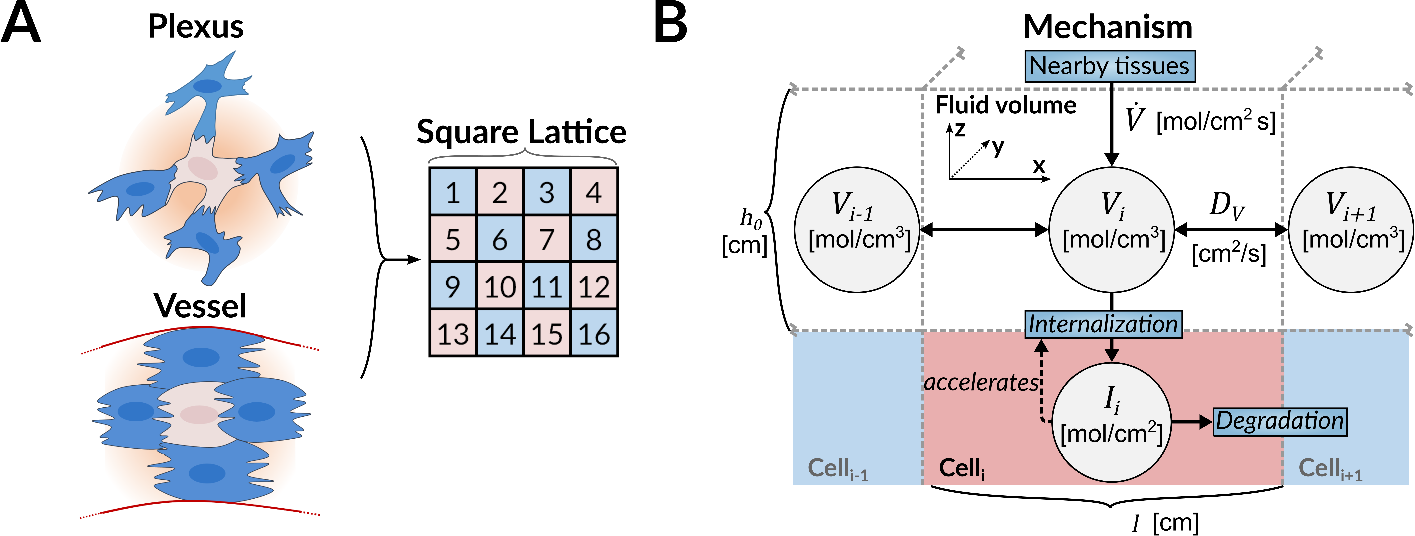
Cellular lattice and reaction-diffusion model. (A) Lattice representation of interconnected endothelial (ECs). Tip cells can emerge in an immature vascular plexus or in a mature vessel; in both cases the structure formed by the interconnected endothelial cells is approximately two-dimensional. We approximate the endothelium as a two-dimensional square lattice, where each square represents an endothelial cell. (B) Reaction-diffusion model on cellular lattice. The diagram presents the idealized representation of the scenario depicted in Fig. 2A. The hypothesis explored here tracks both intracellular (*I*_*i*_) and extracellular (*V*_*i*_) VEGF associated with each cell *i*, so each square in the lattice has both an intracellular compartment and extracellular fluid volume of depth *h*_0_. Extracellular VEGF is introduced from adjacent tissues at a constant rate 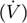 and can diffuse between the fluid volumes above adjacent cells or be internalized into intracellular VEGF. Intracellular VEGF activates signaling processes that accelerate the rate of VEGF internalization in that cell. VEGF is only removed from the system by first-order degradation within the cell.

We translated the hypothetical scenario in Fig. 3B into two ODEs governing the diffusion, internalization, and intracellular signaling of VEGF; a full derivation of the equations is given in Supplemental S2. The concentration of extracellular VEGF in the fluid volume of cell *i* is governed by the following reaction-diffusion equation:

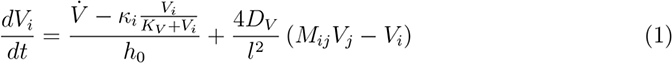

where *κ*_*i*_ [mol/cm^2^ s] refers to the rate that VEGFR2 is internalized into the *i* th cell and *K*_*V*_ [mol/cm^3^] is the dissociation constant of VEGF-VEGFR2 binding. The term *M*_*ij*_*V*_*j*_ represents the average concentration of extracellular VEGF in the fluid volumes of cells *j* that are direct neighbors to cell *i* by employing the connectivity matrix *M*_*ij*_ (38).

The concentration of intracellular VEGF-VEGFR2 is governed by the following differential equation:

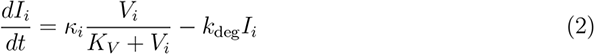

where *k*_deg_ [s^−1^] is the first-order degradation rate of the complexes.

The final component of the diffusive mechanism relates the maximum internalization rate (*κ*) to the concentration of intracellular VEGF-VEGFR2 complexes (*I*). Full activation of certain pathways downstream of VEGFR2 (such as Akt) requires the receptor to be contained in a dynamin-dependent vesicles or endosomes (34, 37), and dynamin activity is under the control of Akt (through eNOS) (31–33). We hypothesized that the movement of VEGFR2 into dynamin-dependent vesicles is the rate-limiting step, as mentioned above, and that the activity of dynamin, and therefore the rate of dynamin-dependent vesicle formation, is governed by intracellular (i.e., in a vesicle or endosome) VEGF-VEGFR2 signaling. We formalized this hypothetical link by making the maximum rate of internalization an activating Hill function of intracellular VEGF-VEGFR2 concentration:

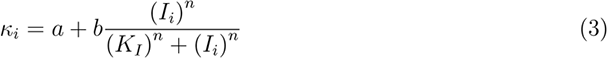

where *a* [mol/cm^2^ s] is the basal internalization rate in the absence of feedback from intracellular VEGF-VEGFR2 signaling, *b* [mol/cm^2^ s] is the gain in the internalization rate that can occur in the presence of strong intracellular VEGF-VEGFR2 signaling, *K*_*I*_ [mol/cm^2^] is the concentration of intracellular VEGF-VEGFR2 complexes when the gain is at 1/2 maximum, and *n* is an effective Hill coefficient for the cumulative signaling and gene regulation processes (Fig. 2B) that connect intracellular VEGF-VEGFR2 signaling to VEGFR2 internalization. In this study, we use a Hill-type representation of the hypothesized signaling processes to provide the necessary nonlinearity for an instability to occur; many regulatory pathways have been successfully modeled with this functional form (20, 24, 39, 40).

We hypothesized that the positive feedback of intracellular VEGF-VEGFR2 complexes on their own creation could cause an instability that enables a cell to have a high rate of VEGF internalization (and therefore a high concentration of intracellular VEGF, strong VEGF-VEGFR2 signaling, and fast VEGFR2 internalization) despite having a low concentration of extracellular VEGF in the local fluid volume. Simultaneously, another cell in the same lattice could have a low rate of VEGF internalization despite having a relatively high concentration of extracellular VEGF in the local fluid volume. When these cells are adjacent to one another, the diffusivity of VEGF enables a continuous flux from the high extracellular (low intracellular) VEGF concentration cell to the low extracellular (high intracellular) VEGF concentration cell, creating a stable pattern. Based on previous literature, we hypothesize that the cells with high levels of intracellular VEGF-VEGFR2 are destined to become tip cells (34–36) and the remaining cells destined to become stalk cells (Fig. 3B, left).

We tested this hypothesis of pattern formation using linear stability analysis and numerical simulation of the non-dimensional form of the governing equations:

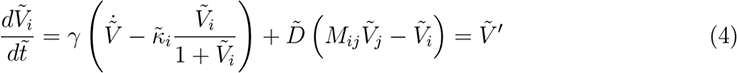

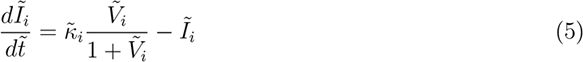

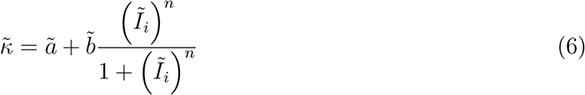

where 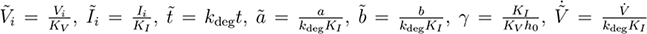, and 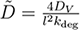.

Before performing the linear stability analysis or simulation, we first identify the uniform steady-states of the governing equations that represent the quiescent endothelium – following stimulation by VEGF but prior to tip cell differentiation. As derived in Supplemental S3), the steady-state values for each species can be calculated as a function of the production rate of VEGF, 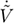:

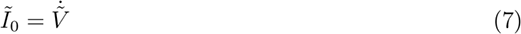

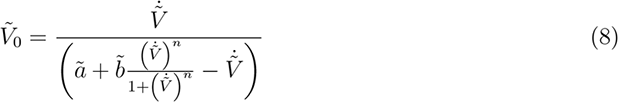

where the subscript 0 denotes the homogeneous steady-state values of each species.

Following Othmer & Scriven (38), the linear stability analysis (detailed in Supplemental S4) transforms the dynamics of nonlinear Eqs. 1 & 2 at the uniform steady-state into a set of *N* (where *N* is the number of cells in the lattice) independent 2 × 2 linear systems of equations. Each independent 2 × 2 system corresponds to a different “structural mode” in the cellular lattice – basic spatial patterns that can grow or shrink over time. As shown in Supplemental S4, these structural modes are represented as the eigenvectors of the connectivity matrix, which have corresponding eigenvalues *q*_*k*_ (that range from −1 to 1 on a square lattice). Considering eigenvectors as a set of wave-like patterns, the structural eigenvalues, *q*_*k*_ are analogous to their wavelengths (see Supplemental Figure S2). For each system of 2 × 2 equations, we calculate a dynamic eigenvalue *λ*_*k*_ (where *k* = 1, 2, …, *N*) that determines the stability of the 2 × 2 system at the uniform steady-state. When *λ*_*k*_ is positive, the system is unstable and deviations from the uniform steady-state on the corresponding structural mode will grow as spatial patterns; when *λ*_*k*_ is negative, the system is stable and deviations from the uniform steady-state on the corresponding structural mode will vanish. The Jacobian matrices we use to calculate *λ*_*k*_ are functions of the structural eigenvalues (*q*_*k*_) of the connectivity matrix, such that each structural mode has differing degrees of stability.

We performed a numerical integration of the governing equations to understand the full impact of the nonlinearity of the mechanism and probe the final steady-states following pattern formation (parameter selection is detailed in Supplemental S5). We primarily looked to the literature to estimate the magnitude of our parameters (provided in Supplemental Table S1). In many cases, we made estimates based on previous computational and experimental studies of VEGF mass transfer and trafficking (41–44). However, where literature estimates were not available, we chose parameters based on the ability of the resulting ODEs to become unstable as predicted by the linear stability analysis. One notable parameter we assign without direct literature precedent is the Hill coefficient (*n*) in our approximation of the positive feedback mechanism in Eq. 3. We assumed a value of *n* = 4; while this number is higher than those used in some previous studies of pattern formation (20, 40), other studies have shown that Hill coefficients generated from numerical fits can be above 3 for single mechanism (45) or effectively greater than 20 for systems of reactions (46).

## Results

### Analysis

We assume that the strength of pro-angiogenic signaling modulates the flux of VEGF towards the endothelium in our model of reaction-diffusion driven tip cell selection (Fig. 2). We sought to understand how the strength of the VEGF flux might impact the formation of tip cell patterns. We analyzed and simulated the mechanism shown in Fig. 3B using the non-dimensional governing equations, Eqs. 4 – 6. Fig. 4A shows the uniform steady-states of VEGF concentration 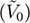 that exist for the given rates of extracellular VEGF production 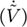, calculated using Eq. 8. Below, we discuss VEGF production rates relative to the production rate where the steady-state VEGF concentration 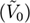 peaks at 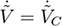.

The segment of the curve in Fig. 4A sloping downwards (where 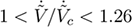) opens up the possibility for an instability: at a uniform initial condition that falls within the negative slope segment, a cell that randomly starts internalizing VEGF faster than its neighbors can sustain that rate of internalization at a lower extracellular concentration of VEGF than neighboring cells; this scenario can occur provided that there is additional flux of VEGF into the fluid volume associated with the cell to balance the increased rate of internalization at steady state. As the cell with faster-than-average internalization depletes the VEGF in its local extracellular volume, a VEGF gradient will begin to emerge that causes VEGF from neighboring volumes to diffuse into its local volume, supplying the necessary additional flux of VEGF.

**Figure 4:**
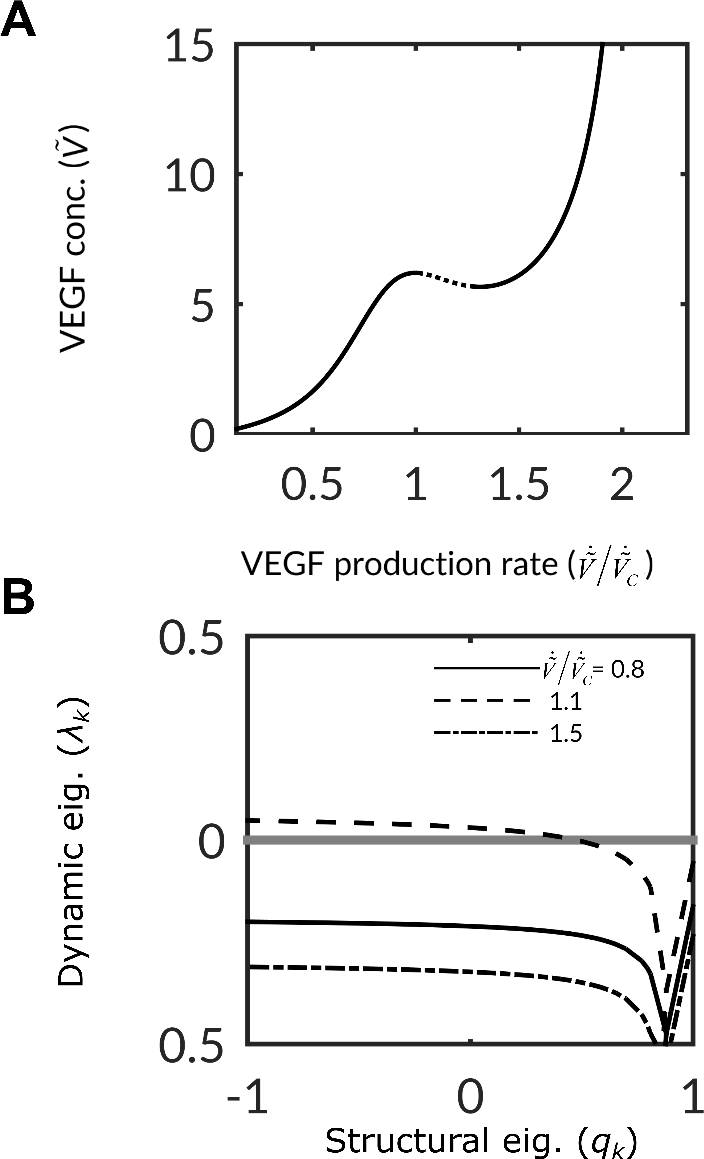
Analysis of uniform steady-states. (A) Uniform steady-states of Eqs. 4–6 as evaluated with Eq. 8 with parameter values in Supplemental Table S1. The x-axis provides the dimensionless production rate of VEGF 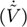; the y-axis provides the dimensionless concentration of VEGF 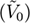 at steady-state. (B) Stability of uniform steady-states. Dynamic eigenvalues (*λ*_*k*_) of the uniform steady-state are plotted against the structural eigenvalues (*q*_*k*_) at various production rates of VEGF 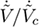. Instability occurs for steady-states with moderate production rates of VEGF – falling on the segment of the curve in (A) with a negative slope.

We first confirmed the existence of this instability by deriving the following stability criterion (full derivation given in Supplemental S4):

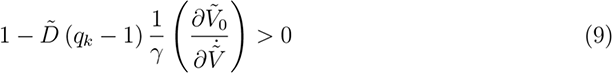

where 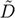 is the dimensionless diffusivity, *γ* is the dimensionless first-order degradation rate of internalized VEGF, 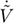 is the dimensionless VEGF production rate, and 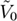 is the dimensionless VEGF concentration at the uniform steady-state. When the inequality in Eq. 9 is satisfied, the uniform steady-state is stable and no patterns of tip cells will emerge from the endothelium; when it is not satisfied, minor deviations in the initial concentration of VEGF will be amplified until a pattern of tip cell selection emerges. The partial derivative in Eq. 9 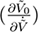 captures the slope of the curve in Fig. 4A. It is clear from Eq. 9 that the partial derivative (and hence slope in Fig. 4A) must be negative for the system to become unstable: the dimensional parameters *γ* and 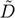 are always positive, and the structural eigenvalue (*q*_*k*_) is always less than 1.

To further analyze the instability, we numerically calculated the dynamic eigenvalues of the uniform steady-state at several representative production rates of VEGF, as shown in Fig. 4B. Steady-states occurring below the critical production rate of VEGF (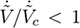 solid curve in Fig. 4B) correspond to a region of Fig. 4A with a positive slope; they are always stable (*λ*_*k*_ < 0). Steady-states which occur at intermediate production rates (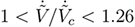; dashed curve in in Fig. 4B) correspond to the region of Fig. 4A with a negative slope; they can become unstable for lower values of *q*_*k*_. Steady-states with higher VEGF production (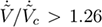; dash-dot curve in Fig. 4B) again correspond to a region with a positive slope; the initial conditions here are also stable of all *λ*_*k*_.

**Figure 5:**
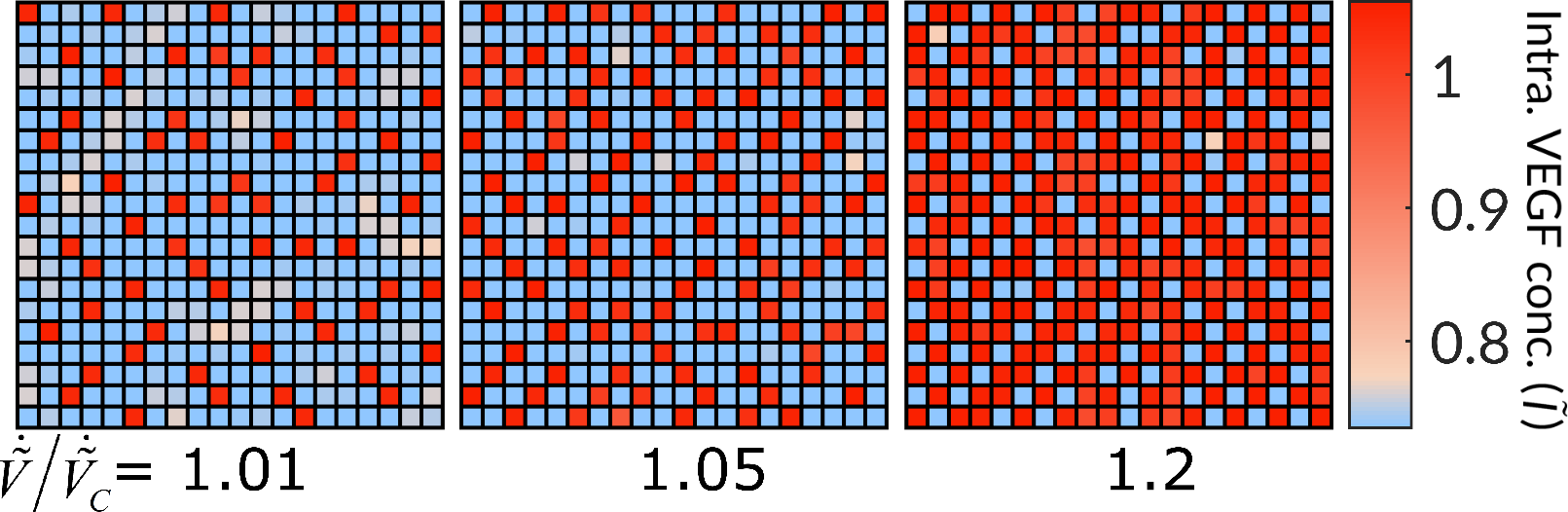
Simulation of tip cell patterning. Intracellular VEGF concentration in the final pattern at various VEGF production rates 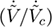. The red cells contain the highest concentration of intracellular VEGF-VEGFR2 complexes, and are therefore the most tip-competent.

### Simulation

We simulated pattern formation under this mechanism for various points along the curve in Fig. 4A. Fig. 5 shows the steady-states that result from simulating pattern formation for several VEGF production rates which slightly exceed the local maximum in Fig. 4A 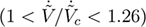. For lower rates of VEGF production (Fig. 5, 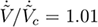), the pattern consists of a small number of tip cells (red) distributed irregularly around the endothelium, with the remaining cells becoming stalk (blue) cells. As the production rate of VEGF increases (Fig. 5, 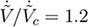), more tip cells emerge in the final endothelium; for high enough VEGF production rates, tip cells outnumber stalk cells in the final steady-state. The location of tip cells within the lattice varies between simulations due to random differences in initial conditions, but the number of tip cells selected remains consistent for a given production rate of VEGF (see Fig. 6B).

Fig. 6A shows the percentage of cells expressing concentrations of intracellular VEGF-VEGFR2 complexes within several ranges in the final steady-states of shown in Fig. 5. The ratio of tip cells 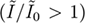 to stalk cells 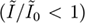 increases with the VEGF production rate, but the concentration of intracellular VEGF-VEGFR2 complexes at either phenotype remains relatively constant.

Fig. 6B shows the number of tip cells counted in the final endothelium as a function of the VEGF production rate. Consistent with the linear stability analysis, distinct tip cells only emerge for moderate VEGF productions rates 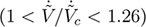. The fraction of tip cells selected jumps discontinuously from 0% to 10% upon increasing beyond 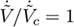, and drops from 84% to 0% as VEGF production increases beyond 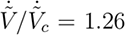. Note that we define there to be 0% tip cells for high-VEGF conditions because we only define tip cells as those having concentrations of intracellular VEGF-VEGFR2 complexes higher than the uniform initial condition; for high production rates of VEGF, all cells will be uniformly high in intracellular VEGF complexes – and under our hypothesis will be expressing tip cell genes – but because the concentrations are equal to the initial condition we do not define “selection” as having taken place. Between the limits of 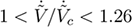, the fraction of tip cells emerging from the endothelium varies continuously and monotonically with respect to the VEGF production rate.

## Discussion

We proposed and analyzed a new hypothesis, depicted in Fig. 2, for the molecular underpinnings of spontaneous endothelial tip cell pattern formation. Our primary goal was to understand if a hypothesis that did not include Notch signaling as the main driver of tip cell selection could exhibit variable density of tip cell selection. The consensus view in the literature is that the density of tip cell selection responds to local stimuli (28, 47) and determines angiogenic sprouting density and microvascular structure (13, 48). Juxtacrine mechanisms based on Notch signaling do not predict this responsiveness they always lead to about 50% of cells becoming tip cells (18, 20). We found that a reaction-diffusion mechanism involving competition among endothelial cells to consume VEGF exhibited a wide range of densities of tip cell selection (see Fig. 5). Below, we will discuss the implications of these results, with an emphasis on interpreting past studies and designing future experiments on endothelial tip cell formation.

**Figure 6:**
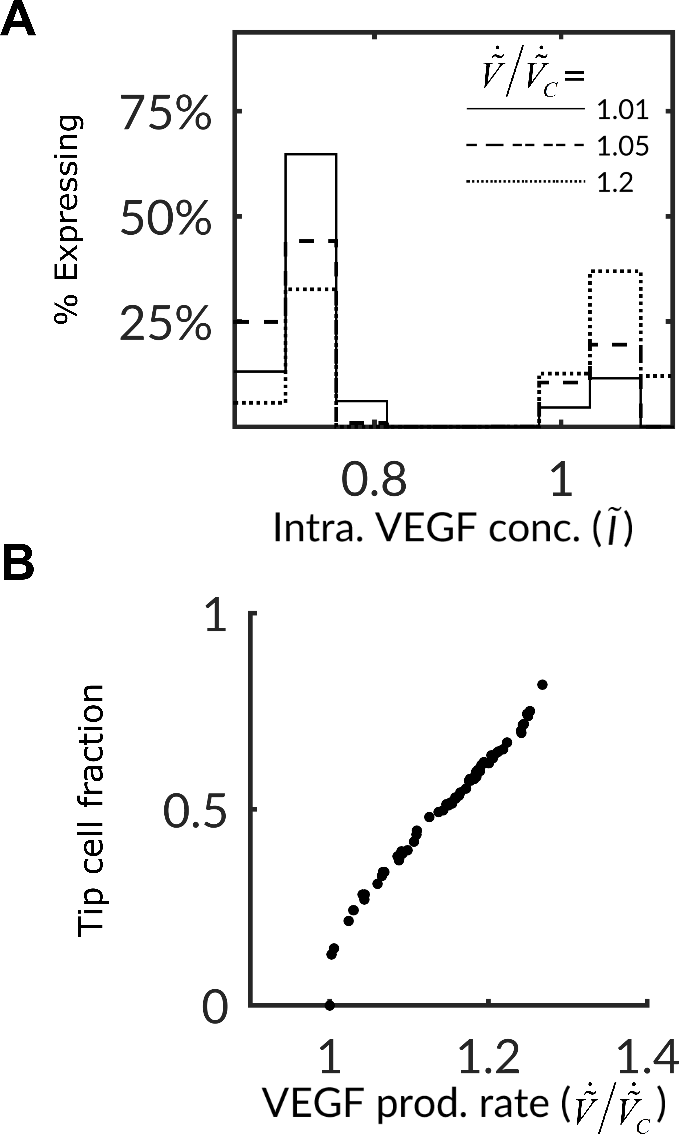
Simulation predictions of expression profile and fraction of tip cells. (A) Histogram of intracellular VEGF concentration 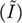 at the final steady-state for three production rates of VEGF 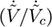. (B) Final fraction of tip cells versus VEGF production rate 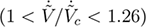. Tip cells were defined as those having a concentration of intracellular VEGF concentrations greater than that of the uniform initial condition 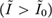 at each VEGF production rate 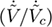. Parameter valsues given in Supplemental Table S1.

A notable feature of this mechanism is the tunability of tip cell selection density with VEGF production rate (Fig. 6B). Research from the early 2000s, oriented towards clinical application of VEGF inhibitors, found that availability of VEGF quantitatively impacted the density (or branching frequency) of the vasculature following angiogenesis (8–11) – including the foundational study on endothelial tip cells (30). However, in the years following, the tip cell selection community focused most experiments on exploring how *Notch* signaling affected tip cell density (12, 16, 47, 49, 50). Only recently has the focus returned to VEGF: a series of experiments showed that tight regulation of the endocytosis of VEGF receptors was essential for proper tip cell formation and sprouting (28, 34–36).

The VEGF reaction-diffusion hypothesis proposed here states that tip cells create a local sink in the concentration of freely-diffusing VEGF in the extracellular space, allowing them to deplete VEGF (29) from their local environment and limit the formation of competing tip cells in their vicinity. Experiments aimed at testing this hypothesis can occur on two fronts: 1) dissecting the hypothetical positive feedback mechanism (shown in Fig. 2B) that allows tip cells to form a local sink in VEGF (proposed here to involve Akt, eNOS, dynamin, and endosomal trafficking), and 2) measuring multicellular outcomes matching the global predictions of the model – such as the continuous dependence of tip cell selection density on VEGF production rate (Fig. 6B). We elaborate on these two proposed paths forward in the following paragraphs.

To dissect the molecular basis of our hypothesis, future studies can test the specific functions we ascribe to the species in our model – namely Akt, eNOS, and dynamin. For example, there are conflicting accounts about whether Akt activation following stimulation of endothelial cells with VEGF is dependent on VEGFR2 internalization: some studies have shown that Akt continues to be activated if full endocytosis of VEGFR2 is prevented (35), but other studies suggest that movement of VEGFR2 from the cell surface into a dynamin-dependent vesicle (not full endocytosis) is the required step before Akt is activated by VEGF-VEGFR2 signaling (34, 37). If it could be shown that inclusion of VEGF-VEGFR2 complexes in dynamin-dependent *vesicles* (and subsequent activation of Akt) is not tightly coupled to VEGF endocytosis and degradation, our proposed mechanism would be incomplete – we would need to identify a signaling pathway that is downstream of VEGF-VEGFR2 signaling from *endosomes* (such as ERK1 (51)) and positively regulates VEGFR2 endocytosis and trafficking to lysosomes (37).

A second avenue for testing our hypothesis is to experimentally examine the validity of our global mechanism – tip cell competition for VEGF through the creation of local sinks (Fig. 3B) – as a whole. We expect that *in vitro* experiments on primary endothelial cells would be the most appropriate method for establishing an empirical relationship resembling the curve in Fig. 4A; this relationship is critical to our biophysical explanation for the pattern-forming instability. Measuring VEGF concentrations *in vivo* is difficult (52), and attempts to eliminate baseline VEGF expression in a whole organism (so that controlled amount of VEGF can be released) are hindered by the embryonic lethality of eliminating even a single allele of VEGF (53). In contrast, previous *in vitro* experiments using endothelial cell monolayers have been successful in measuring the impact of VEGF concentration on cellular responses including gene expression (54), migration (55, 56), and VEGFR2 signaling state (57). The primary challenge in creating an empirical curve like that shown in Fig. 4A is that VEGF flux into the cells 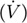 is the imposed parameter in our model, rather than local VEGF concentration; each steady-state VEGF flux has only one possible VEGF concentration, but an endothelial cell can have one of several VEGF fluxes for a given VEGF concentration. Numerous studies have proposed biomaterials that can slowly release VEGF *in vitro* (58); this controlled release could perhaps provide means to impose a steady-state flux of VEGF to a monolayer, but we are unsure how local VEGF concentration at the cell surface might be measured *in vitro*.

We expect that the other important biophysical component of this reaction-diffusion hypothesis – diffusion of VEGF – is better suited to validation through simulation. The transport of a soluble protein like VEGF can be modeled from first principles, and numerous computational studies have predicted how gradients of VEGF can form and be maintained in a variety of contexts, including tip cell formation (41, 59, 60). A three-dimensional simulation of this hypothesis, with accurate models for VEGF diffusion, might be possible once the regulation and kinetics of VEGF endocytosis (i.e., with steady-states like this depicted in Fig. 4A) is elucidated.

The global model prediction of tip cell density being a function of the flux (or production rate) of VEGF might require *in vivo* experiments or tissue engineered *in vitro* constructs which can recapitulate sprouting angiogenesis (6, 61). As mentioned previously, there are limited tools available to measure tip cell selection patterns directly, so using more traditional methods of counting fully-formed tip cells (62) might be a first step towards validating these predictions. One recent study began to test whether the density of tip cell formation responded to changes in VEGF availability and found that spiking the mouse retinal vasculature with concentrated VEGF solutions (1 μg/ml) appeared to eliminate tip cell formation altogether (19). The hypothesis presented here predicts that either adding or subtracting *large* amounts of VEGF could disturb the endothelium to an extent that tip cell pattern formation is no longer functional (see Fig. 6B). In the future, we hope that experiments will be designed around adding or subtracting *small* amounts of VEGF to the endothelium to confirm or refute that incremental changes in VEGF availability have incremental impact on tip cell density.

This study suggests a mechanism for the selection variable tip cell density, a feature missing from previous Notch-based hypotheses of tip cell selection (18, 24). Nonetheless, our model does not explain the myriad previous observations that interfering with Notch has profound effects on endothelial tip cell formation (12, 16, 47). However, our intent is not to disprove that Notch is essential to explaining physiological and pathological sprout formation, but to suggest another biophysical mechanism that the endothelium may use to ensure physiological sprout formation or that tumors may use to attract an aberrantly dense vasculature. Our hope is that future work can reconcile the hypothetical roles of VEGF reaction-diffusion and juxtacrine Notch signaling in coordinating endothelial cell fates as part of a qualitatively new understanding of angiogenesis – giving us a theoretical foothold to much-needed advances in biomedicine and fundamental developmental biology.

## Supporting Material

I. S1 Text: Two-dimensional approximation of diffusion

II. S2 Text: Derivation of governing equations

III. S3 Text: Calculating steady-states

IV. S4 Text: Linear stability analysis

V. S5 Text: Numerical simulation

## Author Contributions

W.B. and A.D.S. developed the hypotheses. W.B. created the governing equations, analyzed the initial conditions, performed the numerical simulation, and analyzed the results. W.B. and A.D.S. wrote the paper.

## Acknowledgments

We would like to thank Jeff Varner and Julius Lucks for the helpful discussions. The work described was supported by the Center on the Physics of Cancer Metabolism through Award Number 1U54CA210184-01 from the National Cancer Institute. The content is solely the responsibility of the authors and does not necessarily represent the official view of the National Cancer Institute or the National Institutes of Health.

## Supporting Citations

Reference (63) appears in the Supporting Material.

